# Label-free droplet image analysis with Cellprofiler

**DOI:** 10.1101/2025.04.04.647160

**Authors:** Dániel Kácsor, Merili Saar-Abroi, Triini Olman, Simona Bartkova, Ott Scheler

**Affiliations:** Tallinn University of Technology

## Abstract

Droplet microfluidic methods used for microbiological experiments are fast, cost-effective, and provide high-throughput data. However, analysis of such image data can be difficult, and detection of molecular labels is limited by microscope parameters.

Currently, there is lack of user-friendly methods to analyse a large volume of label-free droplet images without the need for trained personnel, or expensive, proprietary software. Such methods would make droplet microfluidic technology more widely accessible for a larger range of biological applications.

In this paper we demonstrate an image analysis pipeline designed using Cellprofiler™, a free, open-source software. This pipeline identifies water-in-oil microfluidic droplets, microplastic particles, and bacterial growth without using fluorescent or other labels.

## Introduction

Conventional methods in biology are usually constrained by the availability of resources and time, hindering the scalability and repeatability of experiments. Droplet microfluidics provides a fast, cost-effective way to get thousands of data points from a single experiment (Bartkova et al., 2024). However, analysis of such large datasets often requires elaborate scripting or specific and costly image analysis software. There are currently very few ways to analyze high throughput label-free microscope image data without specific training or prior programming knowledge of the personnel involved (Sanka et al., 2023). This facilitates a need for user-friendly, affordable, and open-source software solutions that can solve this issue (Sanka et al., 2021).

This paper demonstrates an image analysis pipeline which can conduct label-free, brightfield identification of specific objects on 2D microscope images of microfluidic droplets using Cellprofiler™ (Stirling et al., 2021), a free, open-source software. Using this label-free pipeline it is possible to carry out experiments involving the detected objects without using any molecular labels. This not only makes running such experiments easier, but eliminates the chance of possible complications due to toxicity issues or the disruption of cellular processes linked to labels (Alford et al., 2009; Fei & Gu, 2009). Alternatively, it also allows for the design of more complex experiments using other, labelled objects, increasing the total number of detectable objects inside a single microfluidic droplet.

To prove the efficacy of the label-free pipeline, the output from Cellprofiler™ was compared against the true values of objects obtained by manual counting. Images from experiments taken using a confocal microscope have been used, not modified or processed in any way before being analysed inside Cellprofiler™. Figure 1 contains a schematic of the experiment.

**Figure 1:**
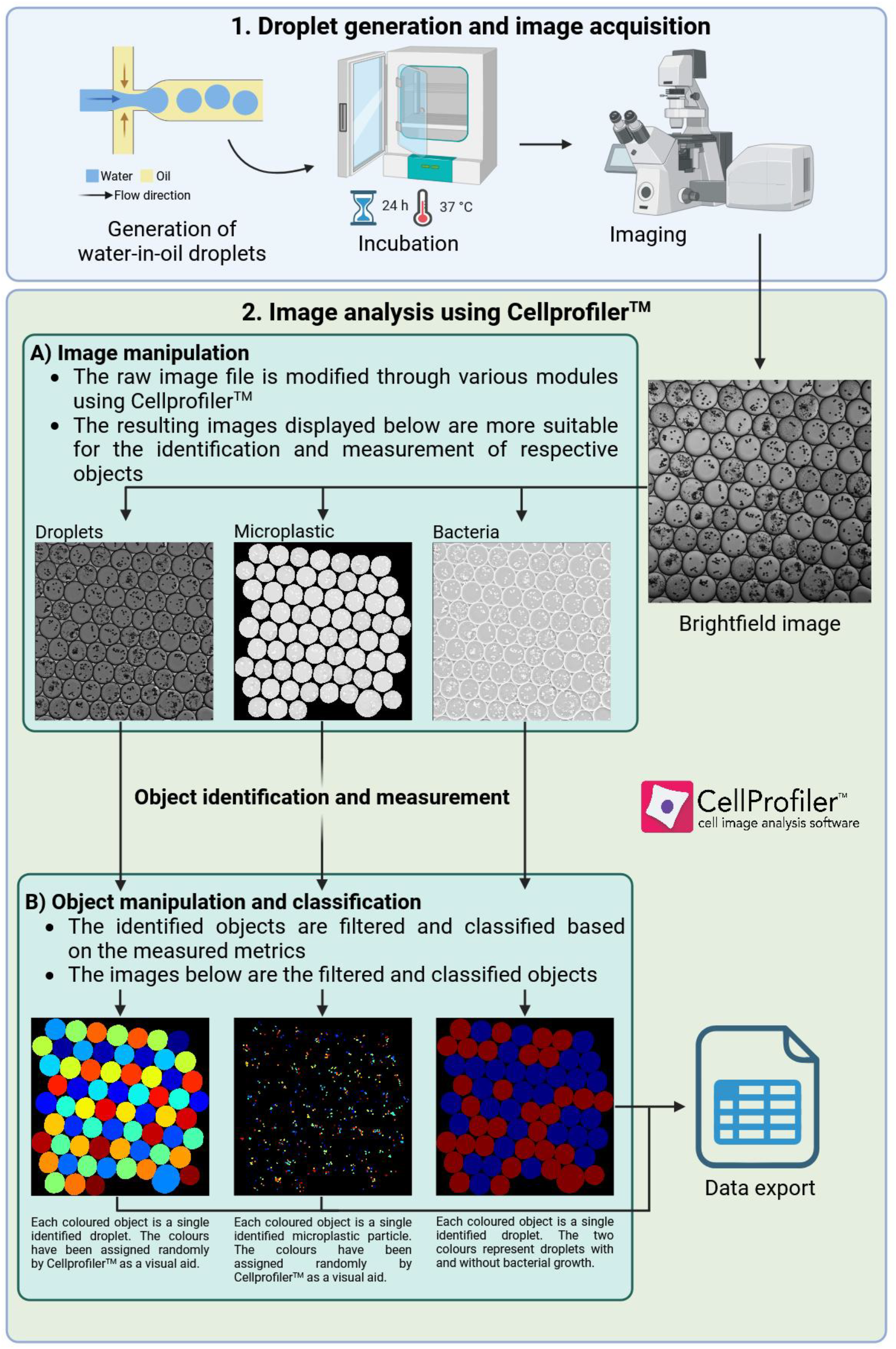
Schematic of the experiment and subsequent image and data analysis. First, monodisperse droplets are generated using liquid samples, further incubated to promote bacterial growth, and imaged under a confocal microscope. The example brightfield image on this figure has been modified for better visibility, and the example bacterial image in 2A is in the penultimate stage of modification, as the final image has very low visibility to the naked eye. The image data then goes through an analysis pipeline in Cellprofiler™. The input image needs to be modified to improve identification accuracy, and the identified objects need to be filtered for good results. Microfluidic droplets and microplastic particles are identified, and droplets are classified based on whether they contain microplastic particles and bacterial growth. The resulting numerical data is exported and assessed using Microsoft Excel and R. Created in BioRender. Kácsor, D. (2025) https://BioRender.com/tyugoor

## Materials and methods

The open-source, script-free image analysis software Cellprofiler™ was used to design and test the label-free image analysis pipeline. Although Cellprofiler™ is primarily designed for analysis of cells, it can be used for a variety of biological applications, including microfluidics, without the need for specialised training (Bartkova et al., 2020). Monodisperse droplets 1,13 nL in volume and 147 µm in diameter on average were generated using a flow focusing PDMS microfluidic chip as described by Bartkova et al. (2020). Samples contained Luria Broth (Biomaxima, Poland) with overnight incubated *Escherichia coli* strain JEK 1036 with a chromosome-incorporated gene encoding the green fluorescent protein (Hol et al., 2015) at 37 °C, and Invitrogen™ Polystyrene CML Latex Beads 10 µM in diameter (Thermo Fisher Scientific, USA). The droplets were imaged using a Countess™ (Invitrogen, Thermo Fisher Scientific Inc., USA) cell counting chamber slide as a monolayer. Roughly 18-20 µL of droplets were pipetted into one chamber. The droplets were imaged using an LSM 900 Laser Scanning Microscope (Zeiss, Germany) with the aid of the software ZEN 3.3 (blue edition). Images with a resolution of 2048×2048 pixels were acquired and exported in TIF format. The images used for analysis are referred to as „brightfield” throughout this paper, however technically they are images taken using differential interference contrast (DIC). As brightfield is more commonly used and easily understood in this field, we have decided to use the term in place of DIC.

The modules and parameters used for the analysis have been chosen based on the software’s documentation, discussion on the online forum forum.image.sc, and empirical, trial-and-error methods. The results can be fully reproduced using the same images and the same modules and parameters. A single example image is provided along with the pipeline for demonstration. In total, 65 images taken from 2 experiments were used for the analysis. 32 of these images were used for the analysis of the total number of plastic particles due to the amount of time manual counting for comparison took. Each image contained 3 types of objects of interest: microfluidic droplets, microplastic particles and bacterial growth.

The “entropy” texture measurement is used for bacterial classification of droplets. Texture measurement was first used for droplet image analysis by Saar-Abroi et al. (2024). A detailed explanation of the pipeline is available in supplementary materials. With 12 workers running at the same time, the heaviest module of the pipeline (CorrectIlluminationCalculate) uses 6 gigabytes of RAM and utilises 100% of an Intel Core Ultra 5 125U CPU.

## Results

The number of identified objects for droplets and microplastic particles, and pairings of droplets with microplastic and bacterial growth were recorded. The results from the analysis using the label-free pipeline were compared against the true values for each type of object, obtained by manual counting. Table 1 contains the resulting data. All data was processed using Microsoft Excel and R 4.4.2 (R Core Team, 2021).

**Table 1:** The table lists the numbers of identified objects for each category. Accuracy is the percentage of correctly identified objects divided by all objects, and precision is the percentage of true positives divided by all positives. True positives and true negatives are correctly identified, false positives are objects that should have been identified as negative, and false negatives are objects that should have been identified as positive. True negatives are not applicable for droplet and microplastic identification. Created in BioRender. Kácsor, D. (2025) https://biorender.com/opb9952

| **Data analysis**<br>**using Microsoft**<br>**Excel and R** | **Number of**<br>**droplets** | **Droplets**<br>**containing**<br>**microplastic** | **Number of**<br>**microplastic**<br>**particles** | **Droplets**<br>**containing**<br>**bacteria** |
| --- | --- | --- | --- | --- |
| **Accuracy** | 96,95% | 99,88% | 83,23% | 100% |
| **Precision** | 99,28% | 100% | 99,16% | 100% |
| **True positive** | 3431 | 3431 | 11245 | 1630 |
| **True negative** | - | 21 | - | 1826 |
| **False positive** | 25 | 0 | 95 | 0 |
| **False negative** | 83 | 4 | 2170 | 0 |

The accuracy of droplet identification is 97%, with precision over 99%. The accuracy of identification of the total number of microplastic particles is 83%, although roughly half of all inaccuracies are due to multiple objects being identified as a single object, and precision is over 99%. This is relevant if we are interested in quantifying the number of particles, however the accuracy of assessing whether a droplet contains any or no plastic at all remains almost 100%, with exactly 100% precision. The accuracy and precision of bacterial identification are exactly 100% each.

Although the pipeline performs well, it has certain limitations. Due to the number and computational strain of new modules necessary for brightfield identification, the label-free pipeline needs up to 4-8 times more time to run, compared to a pipeline using labelled images. Adjustment of certain parameters is also needed to reproduce the accuracy shown in this paper. Specifically, the lower bound of threshold needs to be adjusted for both droplet and plastic identification. This is because the pipeline is sensitive to brightfield image quality, even after illumination correction. This quality is different between microscopes, different image acquisition settings and the quality of the sample itself. Currently, adjustments are needed for each separate sample. While this can be done fast and easily by manually checking 3-4 example images from a given image set, it makes batch analysis of many image sets one after another impossible without a potential drop in accuracy, unless consistent image quality is guaranteed.

## Discussion and conclusions

Our main aim was to create an image analysis pipeline using Cellprofiler that can accurately identify microfluidic droplets, microplastic particles and bacterial growth in brightfield microscope images of monodisperse droplets. The pipeline can identify droplets and microplastic particles with 97% and 83% accuracy, respectively, droplets containing microplastic particles at near 100%, and droplets with bacterial growth at 100% accuracy.

The whole or parts of the pipeline can be used to analyse images from bacterial and microplastic related experiments without the need to use any image analysis software besides Cellprofiler for the entire process. The pipeline takes longer to run compared to one using labelled images, but uses label-free brightfield images without a significant decrease in accuracy. The increased analysis time is due to the increased number and computational demand of modules used, and threshold adjustment is needed due to the difference in image quality between brightfield images.

This pipeline does not require the usage of fluorescent labels, lowering the cost of experiments and reducing the risk of unintended interactions between labels and samples. If needed, it can instead allow for the inclusion of additional labels to increase the number of detectable objects in a single experiment. Using this newly designed label-free pipeline, it will be possible to use microfluidic technology for a wider range of biological applications. It will also increase the accessibility of such technology, as the software’s ease-of-use allows even untrained personnel to learn, operate and modify it for their own use case.

While the pipeline accommodates a wide range of quality for brightfield images, it requires a certain level of quality to be usable. Certain parameters, like <50% object image coverage or foreign objects with a large intensity contrast, can also negatively impact pipeline performance. Although the pipeline is usable in its current state, the need for manual threshold adjustment means bulk analysis of a large number of images without a potential drop in accuracy is only possible if image quality is kept consistent. In the future, the pipeline can be streamlined further to solve the aforementioned issues and expanded to include more objects as needed.

## Supporting information

Explanation of the pipeline

Example image for the pipeline

Image analysis pipeline (zipped)

