## Supplementary material for "Label-free droplet image analysis with Cellprofiler": Explanation of the pipeline

### Detailed description of the image analysis pipeline

Cellprofiler$^{TM}$ is a free, open source software that does not require writing code or the installation of extra plugins. Every module is pre-built and a part of the base installation of the software, only their parameters can be edited.

A) Basic modules which are a necessary part of every pipeline

1. Images: The module responsible for loading image files.
2. Metadata: A module used to optionally extract metadata from image files, which can be used, stored, and exported downstream.
3. NamesAndTypes: A module responsible for assigning a name to image files based on your criteria. All pipelines downstream will use this name to refer to that type of image.
4. Groups: A module used to optionally split image files into groups based on metadata.

B) Modules specific to this particular pipeline

- **Droplet analysis**

5. **CorrectIlluminationCalculate**: The module responsible for illumination correction calculation. One of the most crucial modules used for identifying any type of object on brightfield images. The most important setting is using „each" for function calculation instead of „all", that way the mask is calculated on an image-by-image basis.

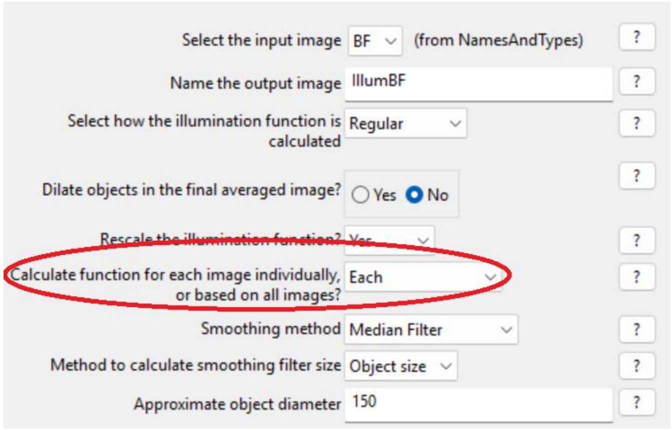


6. EnhanceOrSuppressFeatures: The image is suppressed with a feature size of 5 to smooth out rough edges present due to image resolution and even out intensity

peaks and valleys. Objects are identified more accurately if there are less rough pixelated edges.

**Before** → **After**

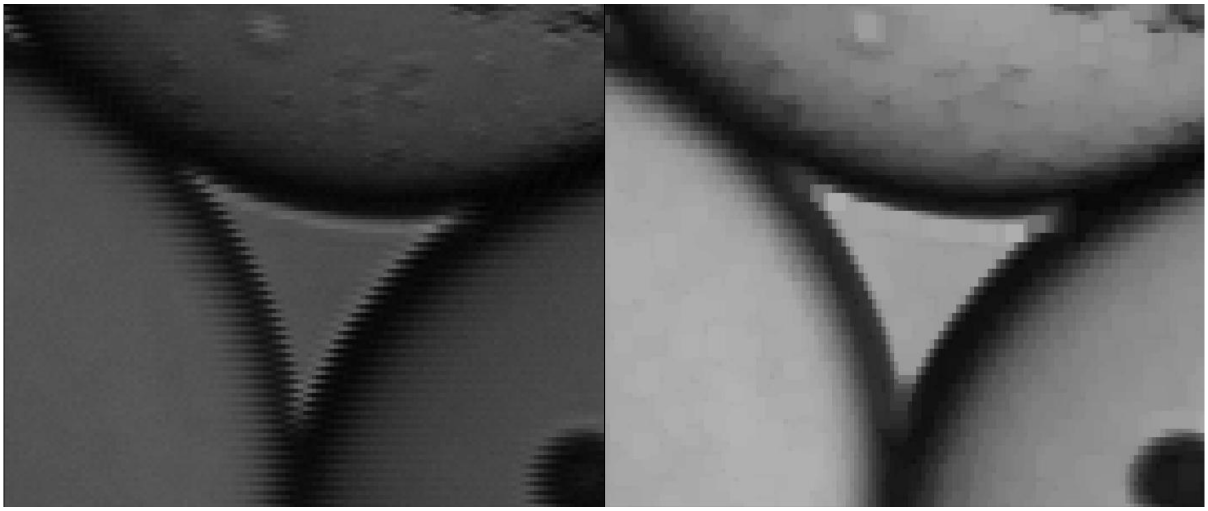

7. CorrectIlluminationApply: The illumination mask calculated before is applied to the image.
8. IdentifyPrimaryObjects: Droplets are identified. The lower bound on threshold parameter needs to be adjusted for good results. Generally, this value should be somewhere between 0.06 and 0.1. The value decides the intensity cutoff during image thresholding, meaning it tells Cellprofiler™ where to draw the line between background and foreground based on the lowest acceptable intensity. If the value is too low, parts of the background bleed into droplets, and if the value is too high, parts of the droplets are cut out, or droplets are split into multiple parts. Sometimes lowering it to the point that some droplets include parts of the background is necessary in order to preserve other droplets on the image. In these cases, all the filtration steps below ensure that these parts are removed and the droplets are restored to their original shapes.

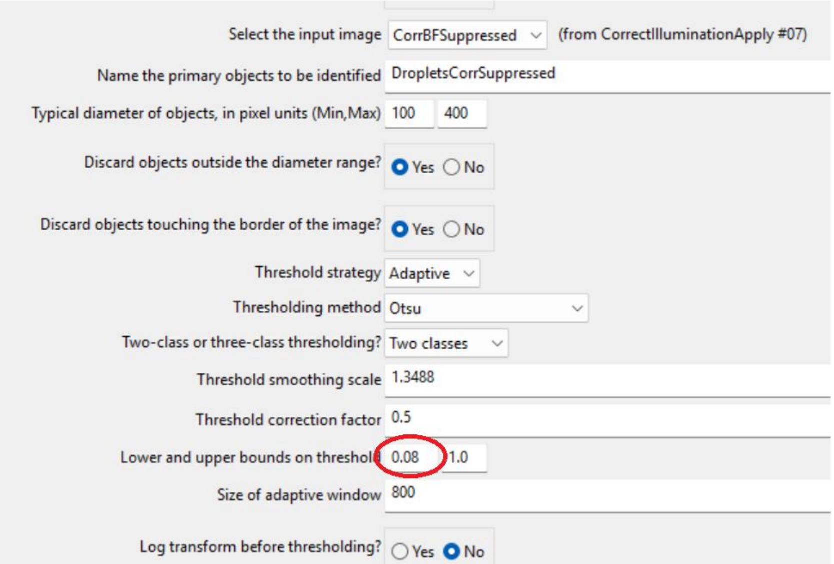


9. MeasureObjectSizeShape: A measurement step. Before objects can be filtered they need to be measured. This means measurement after initial identification and each filtration step.
10. FilterObjects: Objects are filtered by a minimum value of 0.65 for extent and a maximum value of 0.6 for eccentricity. Correct droplets stay in the original image, while droplets with parts from the image background attached, and objects in the image background incorrectly identified as droplets are filtered out and kept as a separate set.
11. MeasureObjectSizeShape: Another measurement step.
12. FilterObjects: A filtration step that aims to separate background objects from droplets with extra parts through the area measurement using a minimum value of 20000. Background objects are generally smaller in size.
13. ErodeObjects: The droplets kepts from the previous filtration are eroded with a feature size of 10. This is usually enough to separate the extra objects from the main body of a droplet.
14. SplitOrMergeObjects: Objects are relabeled, so that the objects separated in the previous step will actually be treated as separate.
15. MeasureObjectSizeShape: Another measurement step.

16. FilterObjects: The relabeled objects are filtered by size with a minimum value of 5000. The extra parts are always much smaller than the droplets. The extra parts are discarded.
17. DilateObjects: Droplets are dilated by 10 to return them to their original size.
18. SplitOrMergeObjects: Droplets are merged just in case some were cut in half by erosion but got kept throughout the process.
19. CombineObjects: Droplets are combined back together with background objects. This is done in case droplets were accidentally filtered out together with the background objects upstream.
20. FillObjects: Objects are filled by convex hull to fill droplets with holes in them. It also makes it easier to filter out background objects downstream.
21. MeasureObjectSizeShape: Another measurement step.
22. FilterObjects: Background objects are ultimately filtered out by form factor with a minimum value of 0.85 and discarded.
23. FillObjects: The original set of correct droplets are also filled by convex hull.
24. CombineObjects: The two droplet sets are combined.
25. MeasureObjectSizeShape: Another measurement step.
26. FilterObjects: Another filtration step for the combined set of droplets, aiming to remove droplets of irregular shape that were not removed so far using form factor with a minimum value of 0.85 and eccentricity with a maximum value of 0.5.
27. DilateObjects: Droplets are dilated with a feature size of 8 to make their size and shape become closer to the true droplet size and shape in the original image.
28. MeasureObjectSizeShape: Another measurement step.
29. FilterObjects: Finally, dilated droplets touching the border are discarded by one last filtration step through the „Image or mask border” filtering mode.

- **Microplastic analysis**

30. ErodeObjects: First, identified droplets are eroded by 8 to exclude droplet edges. If droplet edges are included, parts of them will be falsely identified as plastic particles downstream.
31. CorrectIlluminationApply: The illumination mask we have created before is applied again.
32. MaskImage: The image is masked using the eroded droplets, removing the background and droplet edges.
33. ImageMath: The image is inverted, since Cellprofiler™ recognises bright objects on a dark background more easily.
34. EnhanceOrSuppressFeatures: The image is suppressed with a feature size of 10. While it is somewhat difficult to see any differences on the resulting image with the naked eye, the differences in intensity are perfectly visible to Cellprofiler™.
35. IdentifyPrimaryObjects: Plastic particles are identified. Once again, the lower bound on threshold needs to be adjusted for good results. In this case, the value should generally be somewhere between 0.8 and 1.0. If the value is too low parts of the droplets besides plastic are incorrectly identified as plastic, and if it is too high, many plastic particles stay unidentified.
36. RelateObjects: Plastic particles are paired to droplets in a parent-child object relationship, with plastic as the children.
37. FilterObjects: Identified droplets are filtered by whether or not they have plastic inside, and both sets are kept.

- **Bacterial analysis**

38. CorrectIlluminationApply: The illumination mask is once again applied.
39. EnhanceOrSuppressFeatures: The image is suppressed with a feature size of 3.
40. ImageMath: The image is inverted.
41. EnhanceOrSuppressFeatures: Speckles are enhanced with a feature size of 3, leaving nothing besides bacterial film visible on the image.
42. MeasureTexture: Image texture is measured.
43. ClassifyObjects: Identified droplets are classified based on the entropy texture measurement of the previously modified image, with a cutoff of 2.5. The

measurement type and the value were chosen based on empirical testing. Everything with an entropy measurement below 2.5 gets classified as empty, and everything equal and above as containing bacterial growth.

- Data export

44. ExportToSpreadsheet: This module allows you export your data as a CSV file. As there are many unneeded measurements, it is best to select only the ones needed in order to avoid cluttered files. Any extra data that you want to export needs to be measured first upstream.

Images
Metadata
NamesAndTypes
CorrectIlluminationCalculate
EnhanceOrSuppressFeatures
CorrectIlluminationApply
IdentifyPrimaryObjects
MeasureObjectSizeShape
FilterObjects
MeasureObjectSizeShape
FilterObjects
ErodeObjects
SplitOrMergeObjects
MeasureObjectSizeShape
FilterObjects
DilateObjects
SplitOrMergeObjects
CombineObjects
FillObjects
MeasureObjectSizeShape
FilterObjects
FillObjects
CombineObjects
MeasureObjectSizeShape
FilterObjects
DilateObjects
MeasureObjectSizeShape
FilterObjects
ErodeObjects
CorrectIlluminationApply
MaskImage
ImageMath
EnhanceOrSuppressFeatures
IdentifyPrimaryObjects
RelateObjects
FilterObjects
CorrectIlluminationApply
EnhanceOrSuppressFeatures
ImageMath
EnhanceOrSuppressFeatures
MeasureTexture
ClassifyObjects
**ExportToSpreadsheet**
