## Supplementary figures and images for "Label-free droplet image analysis with Cellprofiler"

### Example image for the pipeline

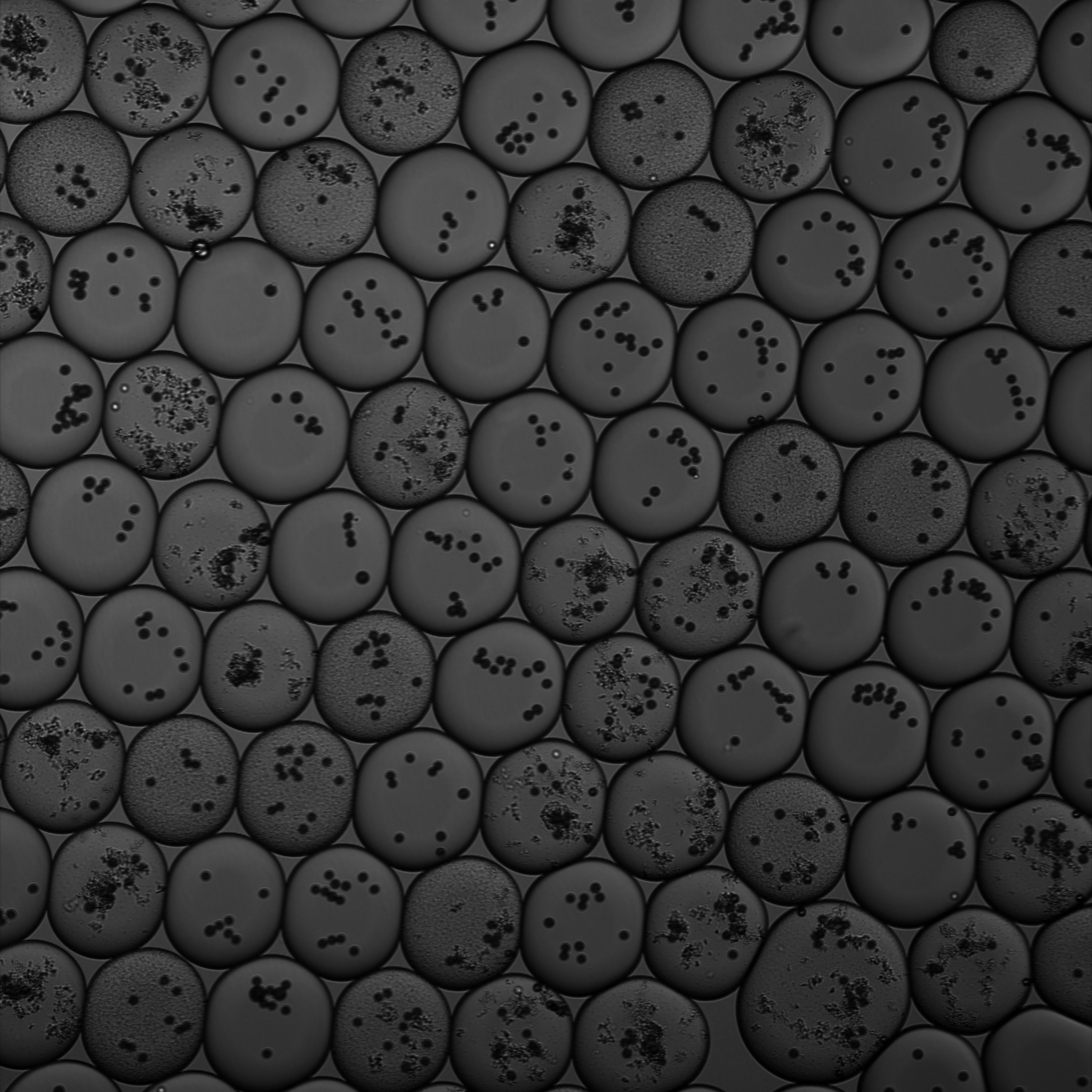
